# Cheaters shape the evolution of phenotypic heterogeneity in *Bacillus subtilis* biofilms

**DOI:** 10.1101/494716

**Authors:** Marivic Martin, Anna Dragoš, Simon B. Otto, Daniel Schäfer, Susanne Brix, Gergely Maróti, Ákos T. Kovács

**Affiliations:** Bacterial Interactions and Evolution Group, Department of Biotechnology and Biomedicine, Technical University of Denmark, Kongens Lyngby, 2800, Denmark; Terrestrial Biofilms Group, Friedrich Schiller University Jena, Jena, 07743, Germany; Disease Systems Immunology Group, Department of Biotechnology and Biomedicine, Technical University of Denmark, Kongens Lyngby, 2800, Denmark; Institute of Plant Biology, Biological Research Center of the Hungarian Academy of Sciences, Szeged, 6726, Hungary

**Keywords:** *Bacillus subtilis*, biofilm, phenotypic heterogeneity, experimental evolution, exopolysaccharide

## Abstract

Biofilms are closely packed cells held and shielded by extracellular matrix composed of structural proteins and exopolysaccharides (EPS). As matrix components are costly to produce and shared within the population, EPS-deficient cells can act as cheaters by gaining benefits from the cooperative nature of EPS producers. Remarkably, genetically programmed EPS producers can also exhibit phenotypic heterogeneity at single cell level. Previous studies have shown that spatial structure of biofilms limits the spread of cheaters, but the long-term influence of cheating on biofilm evolution is not well understood. Here, we examine the influence of EPS non-producers on evolution of matrix production within the populations of EPS producers in a model biofilm-forming bacterium, *Bacillus subtilis*. We discovered that general adaptation to biofilm lifestyle leads to an increase in phenotypical heterogeneity of *eps* expression. Apparently, prolonged exposure to EPS-deficient cheaters, may result in different adaptive strategy, where *eps* expression increases uniformly within the population. We propose a molecular mechanism behind such adaptive strategy and demonstrate how it can benefit the EPS-producers in the presence of cheaters. This study provides additional insights on how biofilms adapt and respond to stress caused by exploitation in long-term scenario.

## INTRODUCTION

Cooperative interactions are prevalent for all life forms [1], even for simple microbes that often exist in communities of matrix bound surface-attached cells called biofilms [2–6]. However, when costly products such as siderophores [7, 8], extracellular polymeric substances [9, 10], digestive enzymes [11], and signaling molecules [12, 13] are secreted and shared, cooperative behavior becomes susceptible to cheating [2, 14–16], where mutants defective in cooperation can still benefit from cooperative community members [4, 5, 17]. It has been shown that spatially structured biofilms, where interactions with clone mates are common and diffusion of public goods is limited, may serve as natural defense against cheating [18–20]. However, long time scale studies have recently reported that biofilm defectors can spontaneously emerge and spread in biofilms by exploiting other matrix-proficient lineages [21–24]. In fact, a pioneering microbial evolution study on *Pseudomonas fluorescens* has already pointed towards dynamic evolutionary interplay between cooperation and exploitation in a biofilm mat [25], where emergence of cellulose overproducer (Wrinkly) allowed mat formation, but also created an opportunity for exploitation by non-producers (Smooth), eventually leading to so called ‘tragedy of the commons’ [4, 26, 27].

Taken together, biofilms are a suitable model to understand social interactions in an evolutionary time scale [23, 28–31]. Modelling and empirical data confirm that mutualism (beneficial to both actor and recipient) and altruism (beneficial to recipient but not to actor) play crucial roles in biofilm enhancement [32] but at the same time can lead to biofilm destabilization [25]. Can cooperators evolve tactics to evade exploitation and in turn, can cheats utilize evolution to enhance their selfish actions?

Recent studies showed that in well-mixed environment, cooperators adapt to cheats by reducing cooperation [14, 15, 33]. Such reduction could be achieved by various strategies, for instance decrease in motility [15], down regulation or minimal production in public goods [14, 15, 33], up-regulation of other alternative public goods [14], or bi-stable expression in virulence gene [2]. Interestingly, populations of cooperators often exhibit phenotypic heterogeneity at the single cell level [34, 35]. Therefore, an alternative and simple mechanism to modulate levels of cooperation in a population would be through changes in phenotypic heterogeneity pattern. However, the long-term effects of cheats on costly goods’ expression at individual cell level, have never been examined. Understanding how heterogeneity of gene expression within the population is affected in the presence of cheats would provide better insight on microbial adaptation and stress response mechanisms.

Here, we address this question using pellicle biofilm model of *Bacillus subtilis* [36, 37]. Pellicle formation in *B. subtilis* involves, amongst others, aerotaxis driven motility and subsequent matrix production [38]. Aerotaxis is important for oxygen sensing to aid cells reach the surface, while matrix formation is significant to sustain cells to adhere to the surface and to each other. Exopolysaccharide (EPS) is a costly public good in *Bacillus subtilis* biofilms [10, 18, 39] and is heterogeneously expressed during biofilm formation with approximately 40% of cells exhibiting the ON state [39, 40]. We aimed to investigate the cheat-dependent alteration related to phenotypic heterogeneity in *eps* expression by the producer.

We reveal that cheating mitigation by the EPS producers involves a shift in phenotypic heterogeneity towards stronger *eps* expression, which can be achieved by a loss-of-function mutation in a single regulatory gene. Our study uncovers an alternative anti-cheating mechanism based on changes in public goods’ expression pattern and highlights meandering trajectories prior cooperation collapse.

## MATERIALS AND METHODS

### Bacterial strains and culture conditions

Strain *B. subtilis* 168 P_hyperspank_-mKATE P_*eps*_-GFP (TB869) was obtained by transforming the laboratory strain, *B. subtilis* 168 P_hyperspank_-mKATE (TB49) [10, 18], with genomic DNA from NRS2243 (*sacA*∷P_*epsA*_-*gfp*(Km)*hag*∷cat) and selecting for Km resistance. The Δ*rsiX* strain with fluorescence reporters (TB959) was obtained by transforming TB869 with genomic DNA isolated from BKE23090 (168 *trpC*2 Δ*rsiX*∷erm) [41]. Strains were maintained in LB medium (Lysogeny Broth (Lennox), Carl Roth, Germany), while 2×SG medium was used for biofilm induction [10]. The Δ*eps* strains (TB608) was created previously [10].

### Experimental evolution

Eight biological replicates of the co-cultures of 1:1 ratio of *B. subtilis* TB869 and TB608 were grown in 48-well plate containing 1ml 2×SG medium at 30°C for two days. Pellicles were harvested into Eppendorf tubes containing 500 μl sterile 2×SG medium and 100 μl of sterile glass sand, vortexed for 90 seconds, 10 μl fraction was transferred into 1ml 2×SG medium of a 48 well plate and incubated at 30°C static condition for two days. Such growth cycle was continuously repeated 35 times. As a control treatment, four biological replicates of mono-cultures of *B. subtilis* TB869 were evolved using the same transfer method. Every 5^th^ transfer (5 growth cycles), harvested cultures were mixed with 15% glycerol and stored at −80°C.

### Population ratio assay

At every 5^th^ transfer, pellicle biofilm productivities and relative frequencies of mutants and WT were qualitatively assessed (colony forming units (CFU)/ml) using LB agar containing selective antibiotics. LB agar plates were incubated at 37°C for 16 h and colonies were counted. Three single clones of WT and of *Δeps* per population per timepoint were isolated from plates and stored at –80°C in the presence of 15% glycerol.

### Pellicle competition assay/Fitness assay

Competition assays were performed as previously described [10]. Specifically, strains of interest were premixed at 1:1 ratio based on their OD_600_ values and the mixture was inoculated into 2×SG medium at 1%. Cultures were grown for 48 h under static conditions at 30°C and their relative frequencies were accessed using CFU counts (and selective antibiotics).

### Stereomicroscopy to assess competition of WT and Δ*rsiX* against Δ*eps*

Fluorescent images of pellicles were obtained with an Axio Zoom V16 stereomicroscope (Carl Zeiss, Jena, Germany) equipped with a Zeiss CL 9000 LED light source and an AxioCam MRm monochrome camera (Carl Zeiss) and HE eGFP (excitation at 470/40 nm and emission at 525/50 nm), and HE mRFP (excitation at 572/25 nm and emission at 629/62 nm) filter sets. Images were taken at 3.5× and 55× magnifications. The exposure times for green and red fluorescence were set up to maximal possible values before reaching overexposure, using range indicator function. Zeiss software was used to obtain overlaid, artificially colored images of both fluorescence channels.

### Qualitative assessment of *eps* expression pattern via laser scanning confocal microscopy

Single isolates of evolved WT (TB869) obtained from population ratio assay were allowed to form 1-day old pellicle. Harvested pellicles were subjected to microscopic analysis using an Axio Observer 780 Laser Scanning Confocal Microscope (Carl Zeiss) equipped with a Plan-Apochromat 63×/1.4 Oil DIC M27 objective, an argon laser for stimulation of fluorescence (excitation at 488 nm for green fluorescence and 561 nm for red fluorescence, with emission at 528/26 nm and 630/32 nm respectively). Zen 2012 Software (Carl Zeiss) and FIJI Image J Software [42] were used for image recording and processing, respectively.

### Flow cytometry and data analysis

Frozen stocks of evolved populations were transferred onto LB-agar plates containing kanamycin (5μg/ml) to select solely for WT colonies. The plates were incubated overnight at 37°C, followed by inoculation of 10 randomly selected single colonies into 2×SG medium. After 24h-incubation at 30°C, the pellicles were harvested, sonicated and diluted accordingly. Flow cytometry was performed using BD FACSCanto II (BD Biosciences). To separate bacterial cells from noise, mKate-fluorescence (constitutively expressed reporter) and GFP-fluorescence (P_*eps*_-GFP promoter fusion) were recorded, gating was setup for mKate-positive objects and GFP signal was measured within these objects. Histograms of P_*eps*_-GFP were created in OriginPro using the same binning intervals for all samples. To remove sample size differences (different amounts of measured objects) histograms were normalized to maximum count, described as Normalized Frequency. To obtained an average distribution image of *eps* expression within populations, a mean count for each histogram bin was calculated (by averaging individual counts within this bin obtained for single isolates), resulting in mean distribution of single cell level *eps* expression per population. Such ‘averaged’ histograms were used solely for visual representation of data and not for statistical analysis.

### Genome re-sequencing and genome analysis

Genomic DNA of single isolates from selected evolved populations were extracted using Bacterial and Yeast Genomic DNA kit (EURx) directly from −80°C stocks grown in LB medium for 5 h at 37°C with shaking at 220 rpm. For population sequencing analysis, approx. 100 colonies belonging to the evolved populations were harvested into 2ml LB broth and incubated at 37°C shaking at 220 rpm for 2-3 h. Re-sequencing was performed on an Illumina NextSeq instrument using V2 sequencing chemistry (2×150 nt). Base-calling was carried out with “bcl2fastq” software (v.2.17.1.14, Illumina). Paired-end reads were further analyzed in CLC Genomics Workbench Tool 9.5.1. Reads were quality-trimmed using an error probability of 0.05 (Q13) as the threshold. Reads that displayed ≥80% similarity to the reference over ≥80% of their read lengths were used in mapping. Quality-based SNP and small In/Del variant calling was carried out requiring ≥10× read coverage with ≥25% variant frequency. Only variants supported by good quality bases (Q ≥ 30) on both strands were considered. Gene functions (product names) in *SI Appendix* Datasets were reported based on SubtiWiki [43].

### Statistical analysis

Statistical differences between two experimental time-points of the same experimental groups (e.g. changes in relative frequencies of WT and Δ*eps* during evolution) were accessed using Pair-Sample t-Test. Statistical differences between two experimental groups were calculated using Two-sample t-Test. To compare multiple samples with WT, we used One-way Repeated Measures ANOVA, and Dunnett Test. ANOVA and Tukey Test was used for multiple samples comparisons.

For analysis of P_*eps*_-GFP expression in evolved populations, we used 10 randomly picked single colonies, cultivated from the frozen stocks (10^th^ transfer) on LB-agar plate with appropriate selection marker (selecting against Δ*eps*). All Flow Cytometry data of P_*eps*_-GFP expression were transferred to histograms, and fitted to Gauss function. Differences in average *eps* expression per population compared to WT_anc_, and differences in single-cell level distribution of *eps* expression compared to WT_anc_ were calculated using One-way Repeated Measures ANOVA, and mean comparison by Dunnett test. Deviation from WT-like distribution were assessed from changes in Adjusted R-Squared (Adj. R. Sq) values for Gauss fitting of Flow Cytometry data. All evolved populations, where average Adj. R. Sq was significantly lower compared to WT ancestor, were suspected to have evolved different phenotypic heterogeneity pattern of *eps* expression. Corresponding histograms were visually inspected, classified as potentially bimodal and subjected to multiple peak fitting. In all such cases, fit quality was improved (Adj. R. Sq>0.98). Mean expression for *eps*-low and *eps*-high subpopulations were compared by ANOVA, Tukey test. No statistical methods were used to predetermine sample size and the experiments were not randomized. One data point was removed from the P_*eps*_-GFP Flow Cytometry dataset of Δ*rsiX* as a significant outlier (P<4.5×10^−7^) confirmed by Grubbs test. All statistical tests and data fitting were performed using OriginPro 2018 software.

## RESULTS

### Cheaters modulate evolution of phenotypic heterogeneity of *eps* expression in the WT

Exopolysaccharide (EPS) is one of the major components of *B. subtilis* biofilm matrix and mutants deficient in EPS production (Δ*eps*) are not able to form pellicle biofilms (Supplementary Fig. 1). In line with previous results [10, 18, 39], we confirmed that the Δ*eps* can take advantage of EPS-producing wild type (WT) and incorporate into the pellicle biofilm, resulting in lower productivity of the WT (Supplementary Fig. 1b) and reduced surface complexity of the pellicle (Supplementary Fig. 1a). Interestingly, despite pellicles formed by the WT+Δ*eps* lacked surface complexity and were more fragile compared to the WT monoculture pellicles (as easily observed during sampling), the total numbers of viable cells (our productivity measure) in the WT and mixed pellicles, were similar (Supplementary Fig. 1b). This indicates high carrying capacity of the WT to support surface colonization by Δ*eps*.

We were further interested if such social cheating could leave a phenotypic or genetic fingerprint in the population of the wild type *B. subtilis*. Previous studies have shown that cooperators can adapt to presence of cheats for example by decreasing the amount of released public goods and therefore minimizing cheating opportunities [2, 14, 15]. As *B. subtilis* exhibits phenotypic heterogeneity in *eps* matrix gene expression [39, 40] (Fig. 1a), we investigated how such heterogeneous expression is influenced by the presence of cheats in an evolutionary perspective. To address this question, we co-cultured the EPS producers (wild type- WT) and cheaters (Δ*eps*) for 10 biofilm growth cycles (~60 generations) starting at 1:1 ratio (see methods). Based on previous studies [21, 44], we assumed that this evolutionary timeframe will be sufficient for evolution of adaptive mechanisms in the WT in response to social cheating, at the same time preventing the diversification of the WT into a biofilm-deficient morphotype, which can be observed on longer evolutionary timescales [21].

**Fig. 1.**
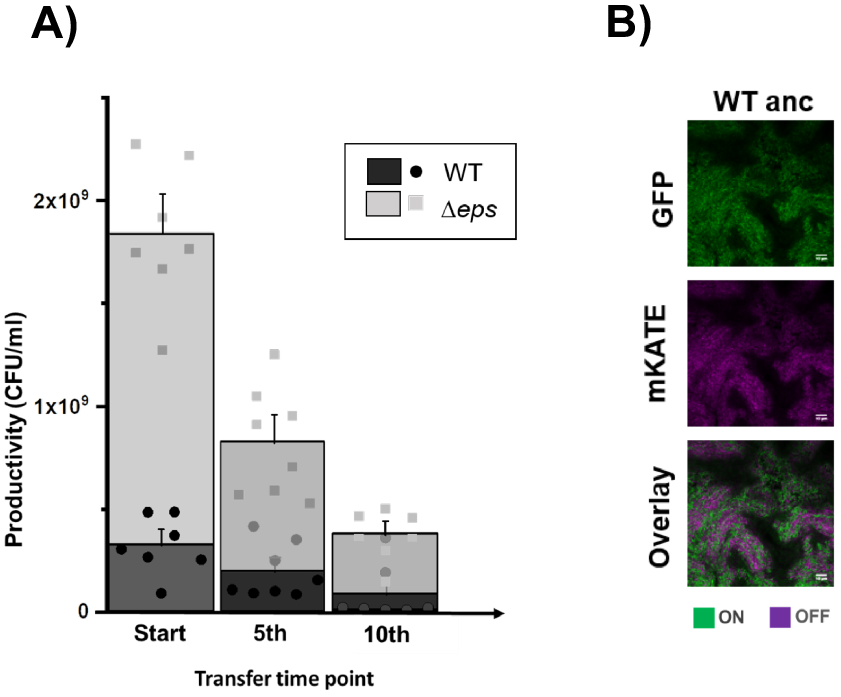
Pellicle productivity and phenotypic heterogeneity. **a** Total colony forming unit per ml of WT and Δ*eps* in 48 h old pellicles non-evolved (Start) (n=9), after experimental evolution at 5^th^ (n=8 populations) and 10^th^ transfer (n=7 populations). One population after 10^th^ transfer was unable to form pellicle attributed to WT being totally outnumbered by Δ*eps*. Dots represent data obtained for all individual populations, while columns represent averages. Error bars correspond to standard error. Levels of Δ*eps* at the start, 5^th^ and 10^th^ transfers were 82%, 76% (p<0.29 compared to start, Pair-Sample t-Test) and 83% (p<0.42 compared to start, Pair-Sample t-Test). **b** Pellicles formed by WT^mKATE^ P_*eps*_-GFP viewed under the confocal laser scanning microscope. Cells constitutively expressing mKATE are represented in magenta (OFF cells) and *ep*s-expressing cells (ON cells) are represented in green. Scale bar 10μm.

During 10-cycle co-cultivation of WT and Δ*eps* strain, we observed a general trend of declining pellicle productivity (Fig. 1b). The relative frequency of EPS non-producers in biofilms was maintained at the high level across all parallel populations (82% at the start, 76% after 5^th^ transfer with p<0.29, and 83% after 10^th^ transfer with p<0.42) (Fig. 1b), indicating that the WT strain was constantly exposed to social cheating throughout the experiment.

Using confocal laser scanning microscopy, qualitative assessment of randomly selected isolates revealed that early populations of the EPS producers (5-10 transfer) exhibited different phenotypes compared to the WT ancestral strain (WT_anc_) (Supplementary Fig. 2). To obtain a quantitative comparison of single-cell level expression of *eps* in the WT_anc_ vs evolved WT populations, we performed flow cytometry measurements of P_*eps*_-GFP harboring strains in pellicles formed by 10 randomly selected isolates per population (Supplementary Fig. 3).

First, we noticed that in most strains isolated from the control evolved populations exhibited an increase in phenotypic heterogeneity in *eps* expression as compared to the WT ancestor (Fig. 2A, Fig. 2B., Supplementary Fig. 3, Supplementary dataset 1). Specifically, while single cell level distribution of P_*eps*_-GFP in the WT ancestor, ideally fitted the Gauss function (Adj. R. Sq = 0.99 ±0.01), this was no more the case for the populations of WT evolved without cheater (C1, C2 and C4; Adj. R. Sq: 0.88-0.93). Similar change was noticed for two WT populations evolved with cheaters (Pop1 and Pop8; Adj. R. Sq: 0.87-0.94). In the aforementioned populations, the *eps* expression was rather bimodal, distributed between low*-eps* and high-*eps* subpopulations (Fig. 2A, Fig. 2B., Supplementary Fig. 3, Supplementary dataset 1). Importantly, these bimodal populations evolved alone (C1, C2 and C4) or coevolved with Δ*eps* (Pop1 and Pop8), were similar in terms of *eps* expression levels or ratios of low*-eps* and high*-eps* subpopulations (Supplementary dataset 1). In addition, an average within-population *eps* expression increased in four out of seven populations that evolved with cheater, but only in one control population evolved without cheats (Fig. 2C., Supplementary dataset 1).

**Fig. 2.**
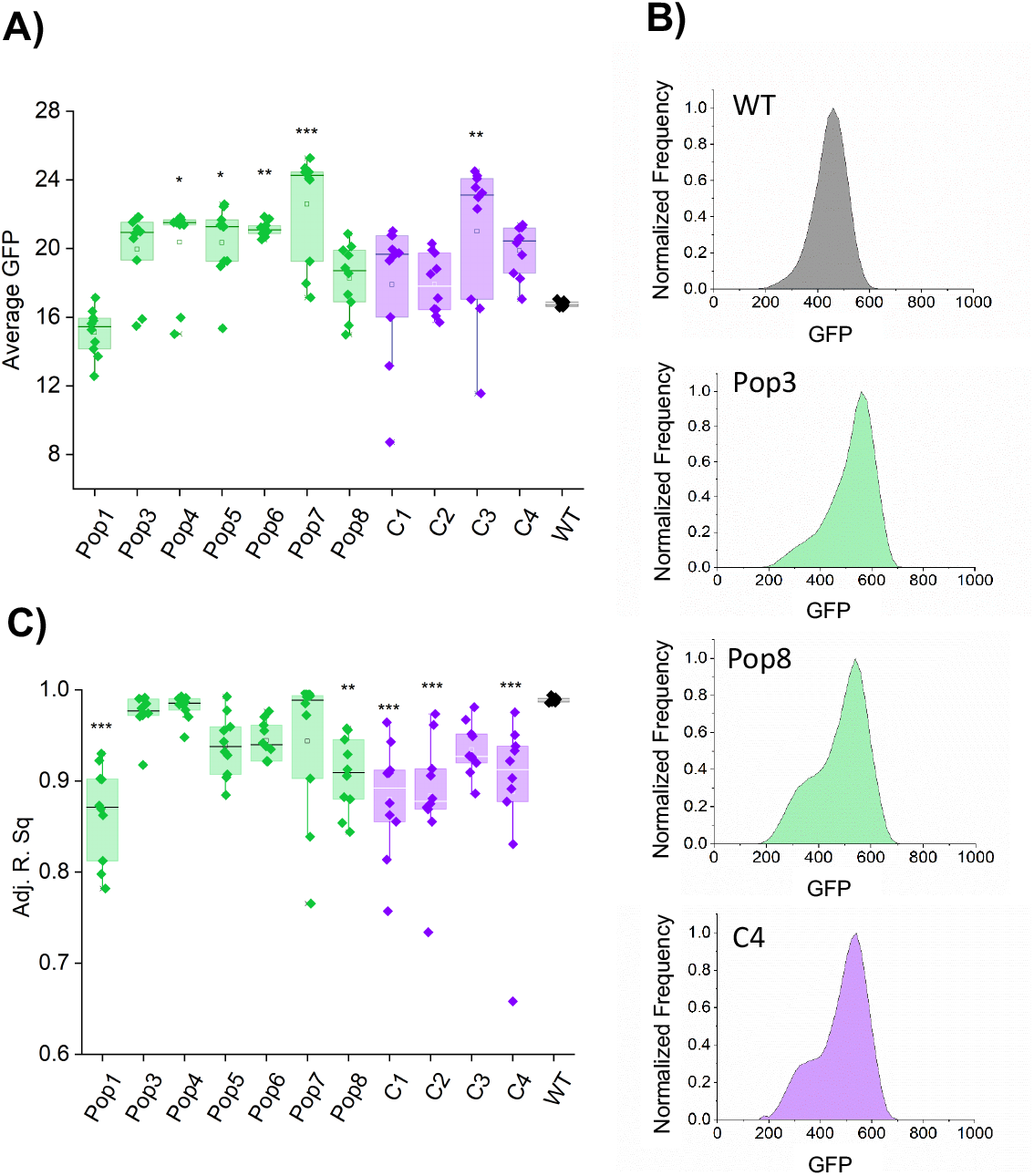
Evolutionary changes in phenotypic heterogeneity pattern and expression of *eps*. **a** Changes in distribution of P_*eps*_-GFP signal in co-evolved and evolved WT populations compared to the WT_anc_ manifested as a decline of adjusted R-square values for Gauss model fitting. **B** Flow cytometry analysis showing average distributions of P_*eps*_-GFP in WT_anc_ (dark grey), WT evolved with cheaters, Pop3 and Pop8 (green) and WT evolved without cheaters, C4 (purple). **c** Average P_*eps*_-GFP expression levels in co-evolved and evolved WT populations compared to the WT_anc,_ calculated from mean values of P_*eps*_-GFP expression within 10 single isolates. For a and c panels *p<0.05; **p<0.01: ***p<0.001 compared to the WT_anc_ (One-way Repeated Measures ANOVA, Dunnett Test). All data and corresponding p values are provided in Supplementary data 1.

Altogether, most populations coevolved with cheater showed an increase in *eps* expression levels, retaining WT-like phenotypic heterogeneity pattern. On the contrary, in majority of control populations (evolved without cheaters) phenotypic heterogeneity level increased, without significant increase in mean *eps* expression (Fig. 2, Supplementary Fig. 3, Supplementary dataset 1).

### Mutations in *rsiX* lead to high-*eps* phenotype

To unravel the genetic basis of the high-*eps* phenotype that evolved in presence of cheats, several single isolates from the evolved populations were subjected to genome resequencing (for details see methods). The comparative analysis of sequencing data revealed that Population 3 and Population 7, co-evolved with Δ*eps*, shared mutations in *rsiX* gene (Supplementary dataset 2). The *rsiX* gene encodes for an anti-sigma factor that controls the activity of extracellular cytoplasmic function (ECF) sigma factor X which is involved in cationic antimicrobial peptide resistance important for cell envelope stress response [45]. Detected mutations resulted either in substitution of Valine 106 to Alanine or frameshift mutations in Serine 104 or Isoleucine 347 that could lead to change or loss of anti-SigX function. Indeed, we were able to recreate the evolved high-*eps* phenotype in the pellicle solely by deleting the *rsiX* gene in the WT ancestor (Fig. 3a,b). Interestingly, a different type of frameshift mutation in Lysine 200 was found in one population of evolved WT alone but this population demonstrated a bimodal phenotypic heterogeneity pattern (Fig. 2, Supplementary Fig. 3, Supplementary dataset 1), suggesting that only certain types of mutations in *rsiX* lead to the uniform shift in *eps* expression or additional mutations have antagonistic effects in this isolate.

**Fig. 3.**
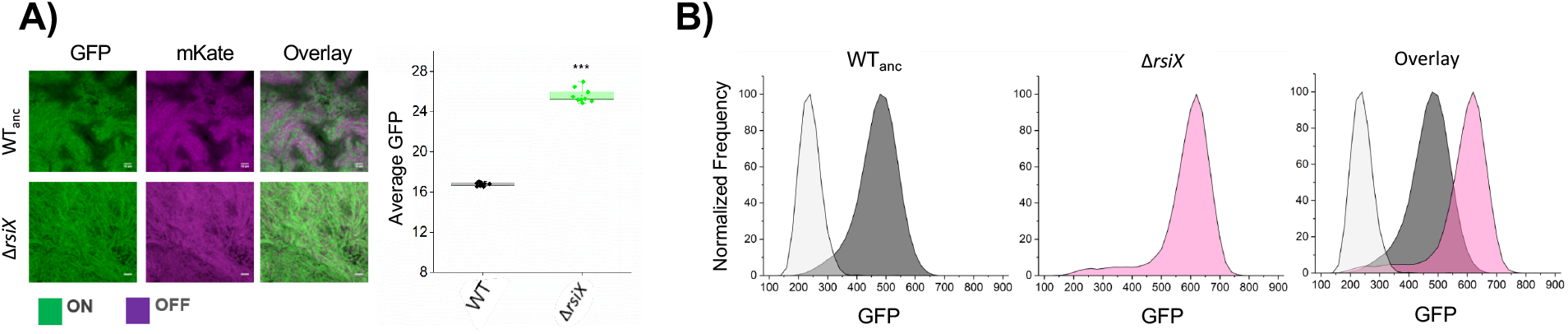
Effect of *rsiX* deletion on *eps* expression in pellicles. **a** Qualitative assessment of *eps* gene expression based on confocal laser scanning microscopy of pellicles formed by Δ*rsiX* showing high-*eps* compared to WT_anc_. Cells constitutively expressing mKATE (OFF) are shown in magenta and *eps*- expressing cells (ON) are represented in green. Scale bar 10μm. Right panel: Average P_*eps*_-GFP expression levels in WT_anc_ and Δ*rsiX* the WT_anc_ calculated from mean values of P_*eps*_-GFP expression within 10 single isolates. ***p<0.001 (Two-Sample t-Test). All data and corresponding P values are provided in Supplementary data 1. **b** Flow cytometry results showing average distribution of fluorescence intensities of WT_anc_ cells (dark grey), Δ*rsiX* cells (pink) and overlay of the 2 in comparison to WT non-labelled (light grey).

### Mutation in *rsiX* contributes to competitive advantage of producer strains against cheats

As mutation in *rsiX* resulted in high-*eps* phenotype that may be linked to elevated secretion of EPS, we hypothesized that Δ*rsiX* producers could support the spread of cheats. To better understand how ancestor WT and Δ*rsiX* interact with Δ*eps*, we cultivated the Δ*eps* in presence of EPS-containing supernatants obtained from the WT and Δ*rsiX* (Supplementary Fig. 4). Both supernatants could partially restore pellicle formation by Δ*eps* resulting in similar productivities of Δ*eps*, thereby not supporting our hypothesis on improved performance of the mutant in presence of high-*eps* Δ*rsiX* strain.

In order to determine the effect of *rsiX* deletion on fitness of the WT in presence of cheats, we performed a series of competition assays. Apparently, the Δ*rsiX* showed two-fold increase in relative frequency (40%) (Fig. 4a, Supplementary Fig. 5) when competed against the Δ*eps*, as compared to the WT ancestor (20%). Additionally, even higher fitness improvement was observed for the WT co-evolved with cheats 5^th^ transfer and 10^th^ transfer, mutually with occurrence of high-*eps* phenotype in those populations. This was not the case for the WT evolved alone at 5^th^ transfer (20%) (Fig. 4a, Supplementary Fig. 5). These results suggest that *rsiX* mutation, which is associated with high-*eps* phenotype, does not fully explain, but contributes to the early improvement of WT competitive strategies against cheats.

**Fig. 4.**
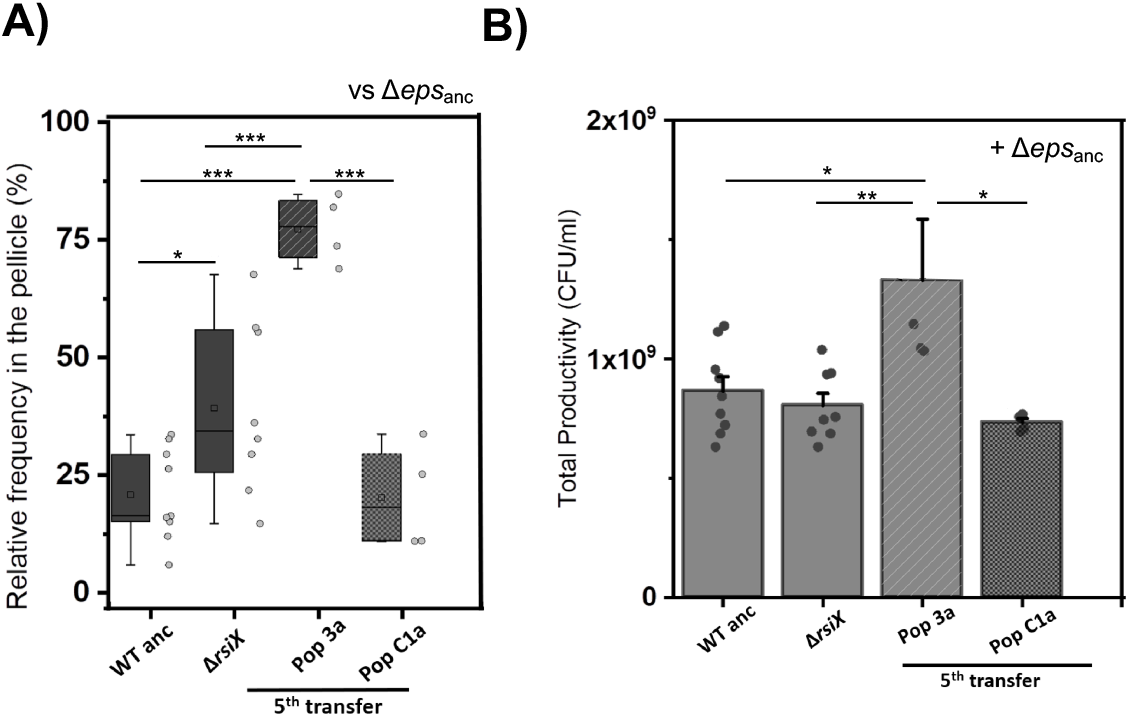
Performance of evolved WT and Δ*rsiX* in co-cultures with Δ*eps*_anc._ **a** Relative frequencies of single isolates belonging to producer populations (WT_anc_ (n=9), Δ*rsiX* (n=8), WT evolved with (n=4) and without cheaters (n=4)) in mixed pellicles with Δ*eps*_anc._ **b** Productivity assay based on total CFU/ml of pellicles of co-cultures of Δ*eps*_anc_ and single isolates belonging to producer populations (WT_anc_ (n=9), Δ*rsiX* (n=8), WT evolved with (n=4) and without cheaters (n=4)). Mean is represented in square within the box plots; median is denoted by horizontal line inside the boxes; whiskers represent the min and max; Error bars in bar graph are based on Standard error; single dots represent the individual data points. For a and b panels: *p<0.05; **p<0.01: ***p<0.001 (ANOVA, Tukey Test).

It is worth to mention that we could not detect any significant fitness costs or benefits linked to *rsiX* deletion in pairwise competition between Δ*rsiX* and WT in the liquid medium (Supplementary Fig. 6; relative fitness of Δ*rsiX* = 1.00 ±0.02 S.D.). Furthermore, we did not observe significant differences in productivities of WT and the Δ*rsiX* mutant, when grown in monoculture pellicles (Fig. 4b), indicating that positive effect of *rsiX* mutation only manifests in presence of cheats. Similarly, different relative frequencies of Δ*eps* in pellicles formed by the ancestor or evolved matrix producers, did not result in different productivities of mixed pellicles (Supplementary Fig. 7). These results suggest that high-*eps* phenotypes are vested on the increase in *eps*-expressing cells or limiting the spread of cheats but do not result in an increase in total yield.

It was previously demonstrated that increased matrix production can allow favorable positioning of a bacterial strain in the biofilm, thereby providing fitness advantage [46]. To test whether high-*eps* phenotype can allow better positioning of the Δ*rsiX* in presence of Δ*eps*, we visualized 48 h grown pellicles formed by Δ*rsiX*:Δ*eps* and WT:Δ*eps* mixtures inoculated at 1:1 initial frequencies (Fig. 5, Supplementary Fig. 8). While WT and Δ*eps* were ‘well-mixed’ with both strains present on the oxygen-rich surface of the pellicle, the Δ*rsiX* strain clearly dominated over the Δ*eps* occupying majority of the biofilm surface and marginalizing the Δ*eps* into small clusters (Fig. 5, Supplementary Fig. 8). Therefore, deletion of *rsiX* and an associated high-*eps* phenotype provides fitness advantage in the presence of Δ*eps* most likely by allowing the EPS producers to occupy upper, oxygen-rich layers of the pellicle. Therefore, *rsiX* frameshift mutation found in certain co-evolved WT populations could be an adaptive mechanism to resist cheating by EPS-deficient strain.

**Fig. 5.**
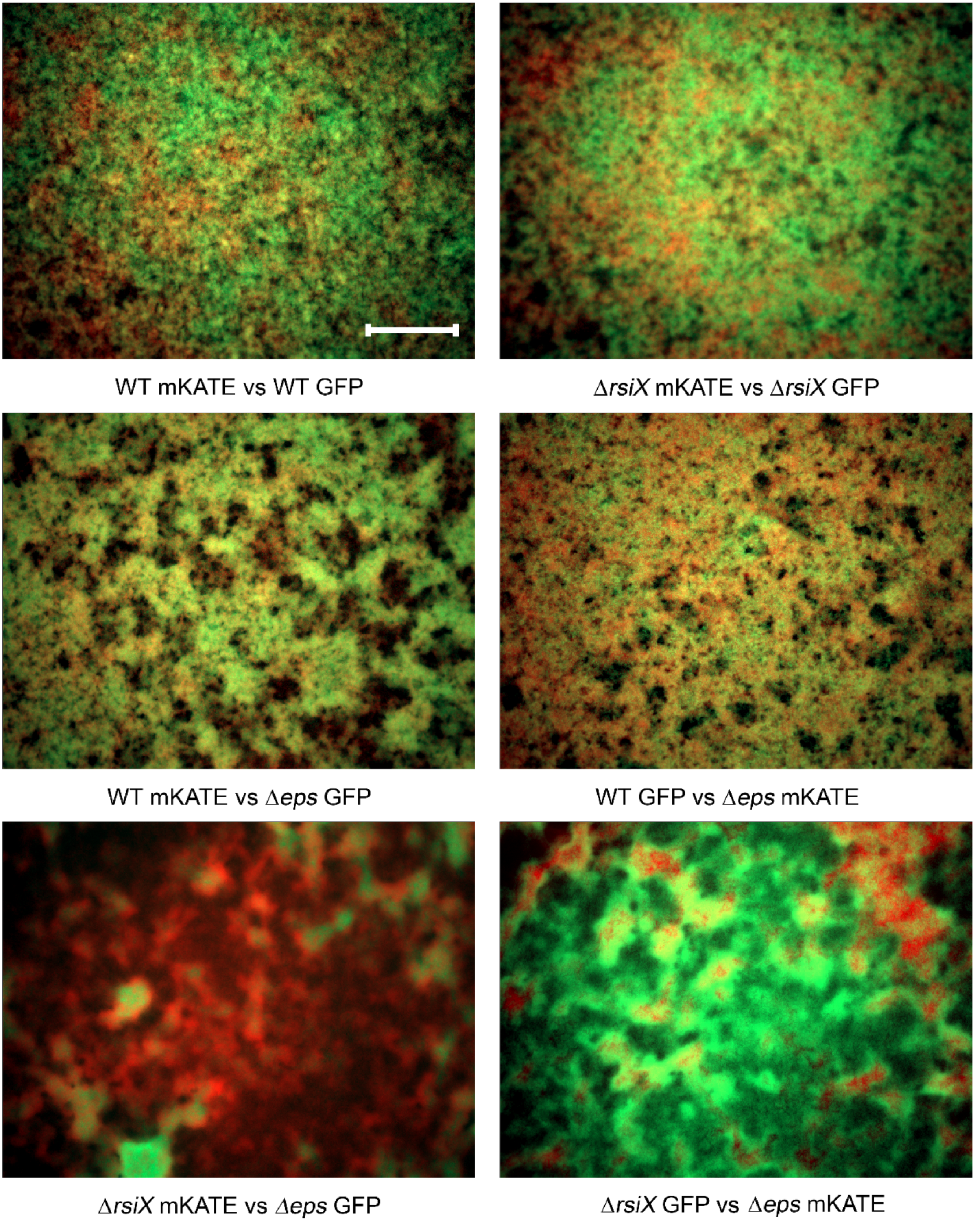
Effect of *rsiX* deletion on positioning of EPS producers in the pellicle. Competition assay between WT+Δ*eps* and Δ*rsiX*+Δ*eps*. Strains labelled with constitutively expressed GFP and mKate proteins, were inoculated in 1:1 initial frequency, pellicles were cultivated for 48 h at 30°C and visualized using stereomicroscope. Upper panels represent controls (two isogenic WT or Δ*rsiX* strains labelled with different fluorescent markers), middle panel represents pellicles formed by WT+Δ*eps* and bottom panels represent pellicles formed by Δ*rsiX*+Δ*eps*, each in two alternative combinations of fluorescent markers. Scale bar corresponds to 500 μm.

### High-*eps* phenotype serves as a transient adaptive response to cheating

As the evolved WT isolates carrying point mutation in *rsiX* as well as the recreated Δ*rsiX* mutant (in ancestral genetic background) performed better in competition with Δ*eps* as compared to the WT_anc_, we reasoned that loss-of-function mutation in *rsiX* together with an associated high-*eps* phenotype, might be an efficient evolutionary strategy against social cheating. Surprisingly however, prolonged evolution experiment eventually led to so called ‘tragedy of the commons’ as the Δ*eps* mutant took over in 6 out of 8 populations, completely abolishing the pellicle formation (Supplementary Fig. 9).

To investigate the genetics behind this phenomenon all evolved WT populations from the last evolutionary time point (or the last time point prior the collapse) were re-sequenced. Curiously, in contrast to the WT populations that were outcompeted by Δ*eps*, both WT populations which resisted the invasion (Pop5 and Pop8), carried mutations in *yvrG* gene (Supplementary Fig. 9, Supplementary Dataset 2) encoding for two-component histidine kinase involved in cell wall process. Finally, the *rsiX* mutation was not detected neither in the last populations before the collapse, with an exemption of population 7, nor at the last transfer point for the non-collapse populations (Supplementary dataset 2), implying that this mutation was lost in the late populations.

## DISCUSSION

Studies on evolution of cooperative behavior is important to understand how social behaviors are shaped in longer time scale. Moreover, exploring long term consequences of exposure to cheating allows to better understand how cooperation prevails in nature where environmental stress and exploitation exist inherently. Here, we took a reductionist approach, focusing on evolution of a single cooperative trait - the expression of *eps*, which plays a crucial role in biofilm lifestyle of *B. subtilis* and other bacteria. As we focused on the single cell level expression of *eps* in multiple single strain, isolated from the ancestral or evolved populations, we could obtain a multi-level insight into evolutionary changes in *eps* expression. Our study revealed previously observed population-level diversification of matrix genes expression, indicating the strain-independence and reproducibility of adaptation in biofilms [21, 24]. Strikingly, next to population-level diversification, we also observe an increase in phenotypic heterogeneity of *eps* expression within single isolates (Fig. 6). Based on co-culture studies performed for WT and Δ*eps* (this work) as well as for WT and spontaneously evolved biofilm-deficient lineage [21], we believe that low-*eps* subpopulations may be acting as conditional cheater, supported by ‘hyper-cooperative’ subpopulations of high-*eps*. It remains to be determined whether increased levels of eps expression translate into higher amount of released EPS, but based on previous studies it is likely to be the case [44].

**Fig. 6.**
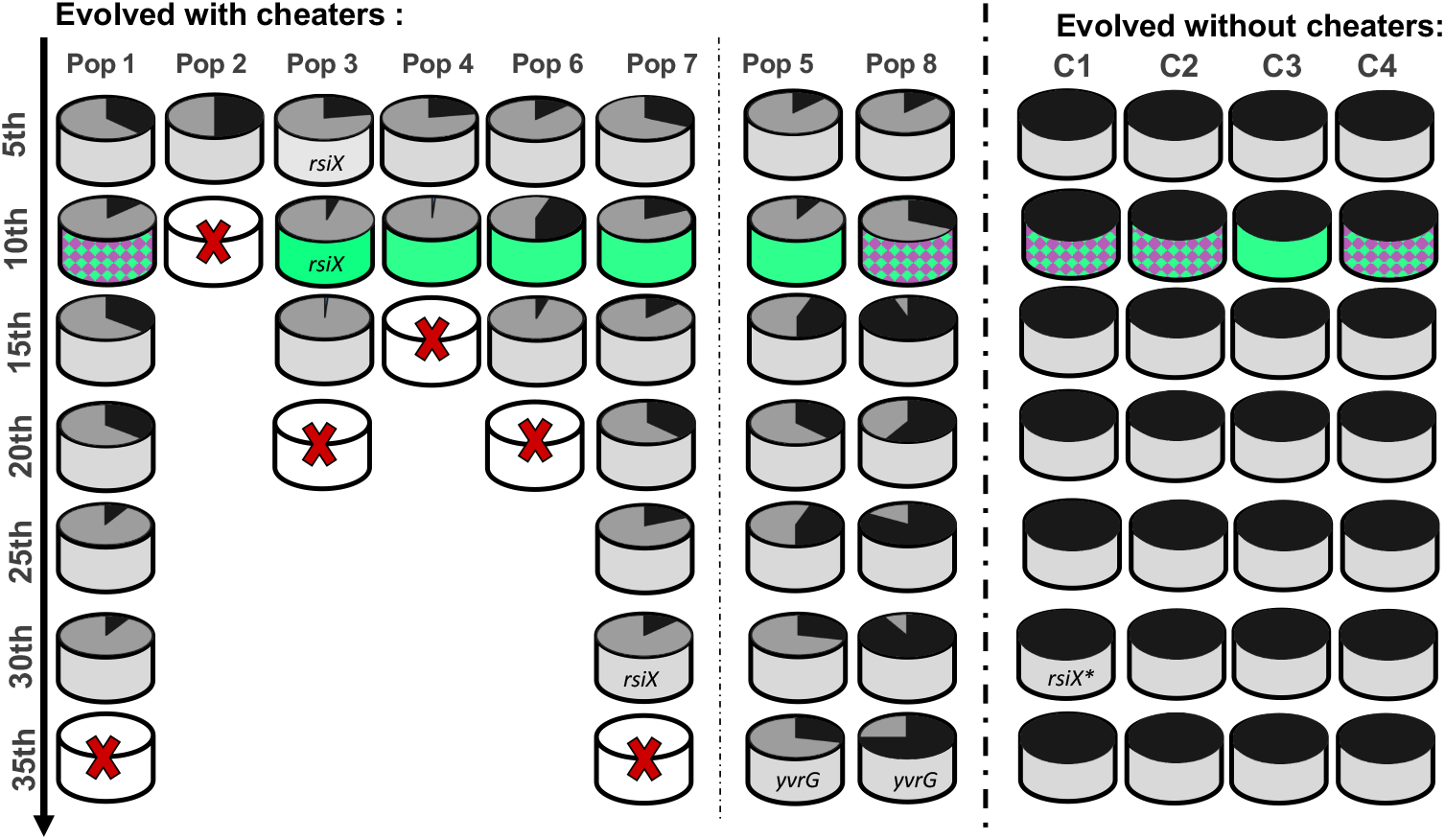
Changes in relative frequencies and *eps* expression pattern during evolution with and without cheaters. Summary figure shows the population dynamics based on producer and cheater frequency per population from 5^th^ transfer to 35^th^ transfers for populations evolved with cheaters with collapse (Pop 1, 2, 3, 4, 6 and 7) and without collapse (Pop 5 and 8) and populations evolved without cheaters (C1, C2, C3, C4) with indications of phenotypes based on high-*eps* or heterogenous *eps*-expression. Key mutations on single clone level of evolved WT and evolved Δ*eps* or population level are specified. The *\*rsiX* mutation differs from mutation observed in strains evolved with cheaters.

Previous evolution studies on cheater-cooperator interactions in spatially structured environment showed cheater mitigation via minimization of the cooperative trait [2, 14, 15]. On the contrary, here we show that cooperators respond to cheating by intensifying the cooperative behaviors through uniform shift towards higher *eps* expression. Further molecular analysis of the high-*eps* isolates strongly suggests that this phenotype is triggered by loss-of-function mutation in *rsiX* gene. The product of *rsiX* represses the activity of ECF sigma factor, SigX that is involved in cell envelope stress response against cationic antimicrobial peptides [47]. Importantly, SigX has been previously shown to induce expression of *epsA-O* in *B. subtilis* via a complex regulatory pathway involving Abh and SlrR [48], explaining the observed enhanced in *eps* gene expression in *rsiX* mutant. Another example of matrix overproduction via ECF adaptation was also reported in Gram-negative bacterium *Pseudomonas aeruginosa* where mutations in another ECF called AlgT led to alginate overproduction and increased resistance to antimicrobials [49]. Therefore, adaptive boosts in matrix production through modulation of ECF is not exclusive for *B. subtilis* but seems to occur also in medically relevant Gram-negative pathogens like *P. aeruginosa*.

In contrast to previous studies that addressed long term cheating on diffusible siderophores [50–53], we explored evolutionary interplay between biofilm producers and non-producers in structured environment. Our results support previous observations on evolution of specific cheating-resisting mechanisms in co-operators, pointing towards ubiquity of this phenomenon. In addition, our work brings up two major findings 1) matrix producers can adapt to matrix non-producers by shifting phenotypic heterogeneity towards increased levels of matrix-expression, 2) high-*eps* phenotype is associated with favorable positioning of the matrix producers in the biofilm in presence of cheats, thereby limiting their numbers, 3) high-*eps* anti-cheating strategy is a short-term solution followed by tragedy of the commons. As EPS-deficient strain took over in all but two mixed populations (including populations, without *rsiX* mutation and homogenous shift towards higher *eps* expression), we do not interpret the collapse as a direct consequence of mutation in *rsiX* gene. However, we argue that an emergence of several matrix overproducing lineages, may facilitate the spread of cheats [21], especially if a substantial number of cells within the high-*eps* lineage serves as facultative (phenotypic) cheaters. As recently demonstrated, EPS-deficiency is not a dead-end strategy for *B. subtilis* population, because alternative EPS-independent biofilm formation strategies can emerge by single amino acid change is TasA [44]. It remains to be discovered whether shifts in phenotypic heterogeneity in response to long term cheating is general phenomenon that applies to different types of public goods.

## Acknowledgements

The authors thank James Gurney for suggestions.

This work was funded by the Deutsche Forschungsgemeinschaft (DFG) to Á.T.K. (KO4741/2.1) within the Priority Program SPP1617. M.M. was supported a FEMS Research and Training Grant (FEMS-RG-2017-0054). This project has received funding from the European Union’s Horizon 2020 research and innovation programme under the Marie Skłodowska-Curie grant agreement No 713683 (H.C. Ørsted COFUND to A.D.). Work in the laboratory of Á.T.K. is partly supported by the Danish National Research Foundation (DNRF137) for the Center for Microbial Secondary Metabolites.

## Competing Interests

The authors declare that there are no competing financial interests in relation to the work described.

## Authors contributions

Á.T.K. conceived the project, M.M., A.D., S.B. and D.S. performed the experiments. G.M contributed with methods. M.M., A.D. and Á.T.K. wrote the manuscript, with all authors contributing to the final version.

## Supplementary Figures and Dataset

**Fig S1. Pellicle biofilm formation and total productivity assessment.**

**a)** Pellicle biofilms formed by mono-cultures of WT, Δ*eps*, and co-culture of WT+ Δ*eps* in 2xSG medium incubated for 48 hours at 30°C recorded using Samsung Galaxy S6 Phone Camera. Scale bar, 1cm. **b)** Productivity assessment based on CFU/ml were performed on pellicle biofilms of WT, Δ*eps*, WT+ Δ*eps* co-culture and control co-culture of two WT strains (WT^mKate^ used in the evolution experiment, and non-labelled WT). Productivity of Δ*eps* dramatically increased (p<3×10^−5^, Pair-Sample t-Test) when co-cultured with WT, while productivity of WT decreased (p<0.002, Pair-Sample t-Test) in the presence of Δ*eps*, indicating the ability of the mutants to act as cheaters (n=5, error bar based on standard error).

**Fig S2. Qualitative assessment of *eps* gene expression based on confocal laser scanning microscopy.**

Pellicles formed by randomly selected WT strains 168 mKATE P_*eps*_-GFP evolved in the presence of cheaters (Pop1-8) and in the absence of cheaters (C1-C4) were visualized using confocal laser scanning microscope. Cells constitutively expressing mKATE are represented in magenta (OFF cells) and *ep*s- expressing cells (ON cells) are represented in green. Scale bar 10μm.

**Fig S3. Single cell level distribution of *eps* expression in 10 randomly selected single isolates from WT_anc_ and evolved WT populations.** Flow low cytometry data (BD Facscanto II, BD biosciences) showing single cell level distributions of fluorescence intensity of 24-hour old pellicles established by 10 randomly selected single isolates from populations of WT evolved in the absence (C1-C4) or presence of cheaters (Pop1-8) as well as autofluorescence distribution in non-labelled WT control.

**Fig S4. Complementation assay of Δ*eps* with supernatant from Δ*rsiX* or WT.** Productivity data of pellicles produced by the complementation showed that hyper ON Δ*rsiX* mutant does not contribute to improved performance of Δ*eps*. Mean is represented in square inside the box plots; median is denoted by horizontal line within the boxes; whiskers represent the min and max (n=3 for Δ*eps*; n=6 for Δ*eps*+SMs). ***p<0.001 compared to Δ*eps;* Δ*eps*+ WT SM is not significantly different from Δ*eps*+ Δ*rsiX* SM (p=0.58) (ANOVA, Tukey Test).

**Fig S5. Performance of Δ*rsiX* and evolved and co-evolved WT in mixed pellicles with Δ*eps*_anc_.** Pellicle competition assay of single clones belonging to producer populations (WT_anc_ (n=9), Δ*rsiX* (n=8), WT evolved with (n=4) and without cheaters (n=4)) against Δ*eps*anc. Mean is represented in square inside the box plots; median is denoted by horizontal line within the boxes; whiskers represent the min and max; single dots represent the individual data points (n). ***p<0.001 compared to the WT_anc_ (One-way Repeated Measures ANOVA, Dunnett Test).

**Fig S6. Fitness effects of *rsiX* deletion.** Selection rate based on fitness assay in pairwise competition of Δ*rsiX* and WT ancestor showing no significant fitness cost brought about by *rsiX* mutation. Relative fitness of Δ*rsiX* is 1.00 ± 0.024 SD. Mean is represented in square within the boxplots; median is denoted by horizontal line inside the boxes; whiskers represent the min and max.

**Fig S7. Pellicle productivity of monocultures.** Total CFU/ml of pellicles produced by mono-cultures of WT_anc_ (n=7), Δ*rsiX* (n=3) and evolved with cheaters (n=8) from population 3 (5^th^, 10^th^) and 7 (30^th^) and single clones of WT evolved without cheaters (n=4). Mean is represented in square; median is denoted by horizontal line inside the box; whiskers represent the min and max; single dots represent the individual datapoints (n). All p values >0.05 (ANOVA, Tukey Test).

**Fig S8. Effect of *rsiX* mutation on positioning in the pellicle.** Competition assay between fluorescently WT+Δ*eps* and Δ*rsiX*+Δ*eps*. Strains labelled with constitutively expressed GFP and mKate proteins, were inoculated in 1:1 initial frequencies, pellicles were cultivated for 48h at 30⁰C and visualized using stereomicroscope. Upper panels represent controls (two isogenic WT or ΔrsiX strains labelled with different fluorescent markers), middle panel represents pellicles formed by WT+Δ*eps* and bottom panels represent pellicles formed by Δ*rsiX*+Δ*eps*, each in two alternative combinations of fluorescent markers. Well size = 1.5cm.

**Figure S9. Productivity changes during short and long-term co-evolution of WT+Δ*eps*. Pellicle** Total colony forming unit per ml of WT and Δ*eps* in 48-hour old pellicles **A)** non-evolved (start) (n=9), after experimental evolution at 5^th^ (n=8 populations) and 10^th^ transfer (n=7 populations) One population after 10^th^ transfer was incapable to form pellicle attributed to WT being outnumbered by Δ*eps*. **B)** Line graph showing the fate of populations. Data was obtained from CFU assay using selective antibiotic marker, Kanamycin selecting for WT and Tetracycline for selecting Δ*eps*.

**Supplementary dataset 1.** Statistical comparisons of *eps* expression and single cell level distribution within ancestral and evolved populations.

**Supplementary dataset 2.** List of detected SNPs of evolved strains and their functional analysis.

## Notes

#### Summary of Updates

Manuscript has been revised and substantially expanded based on reviewers comments after new rejection from another journal.

