## Supplementary Figures S1 to S9 for "Cheaters shape the evolution of phenotypic heterogeneity in *Bacillus subtilis* biofilms"

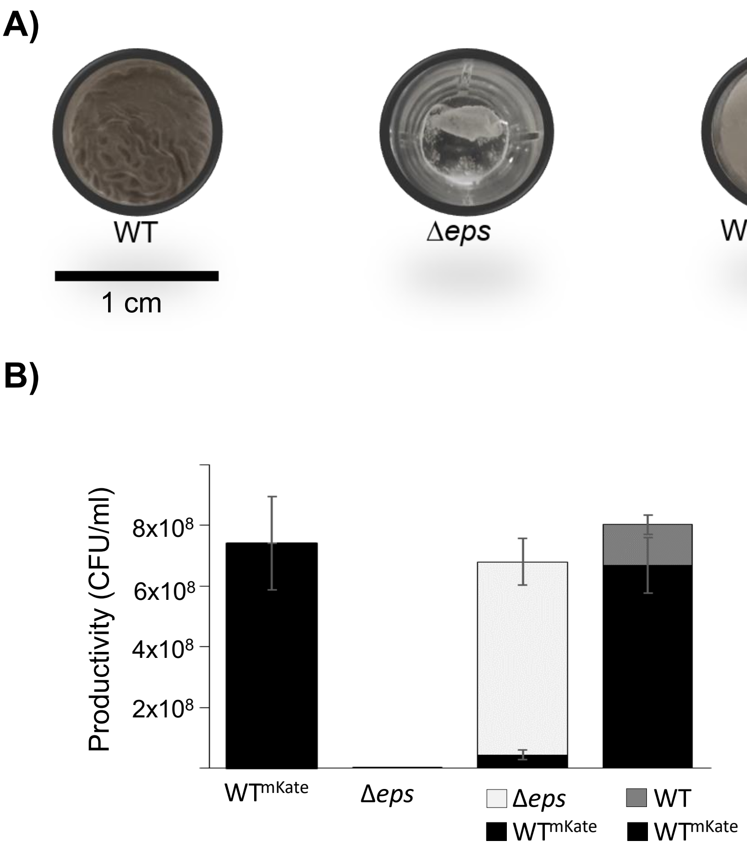


**Fig S1. Pellicle biofilm formation and total productivity assessment. a)** Pellicle biofilms formed by mono-cultures of WT, Δ*eps,* and co-culture of WT+ Δ*eps* in 2xSG medium incubated for 48 hours at 30°C recorded using Samsung Galaxy S6 Phone Camera. Scale bar, 1cm. **b)** Productivity assessment based on CFU/ml were performed on pellicle biofilms of WT, Δ*eps,* WT+ Δ*eps* co-culture and control co-culture of two WT strains ($WT^{mKate}$ used in the evolution experiment, and non-labelled WT). Productivity of Δ*eps* dramatically increased ($p<3x10^{-5}$, Pair-Sample t-Test) when co-cultured with WT, while productivity of WT decreased ($p<0.002$, Pair-Sample t-Test) in the presence of Δ*eps*, indicating the ability of the mutants to act as cheaters (n=5, error bar based on standard error).

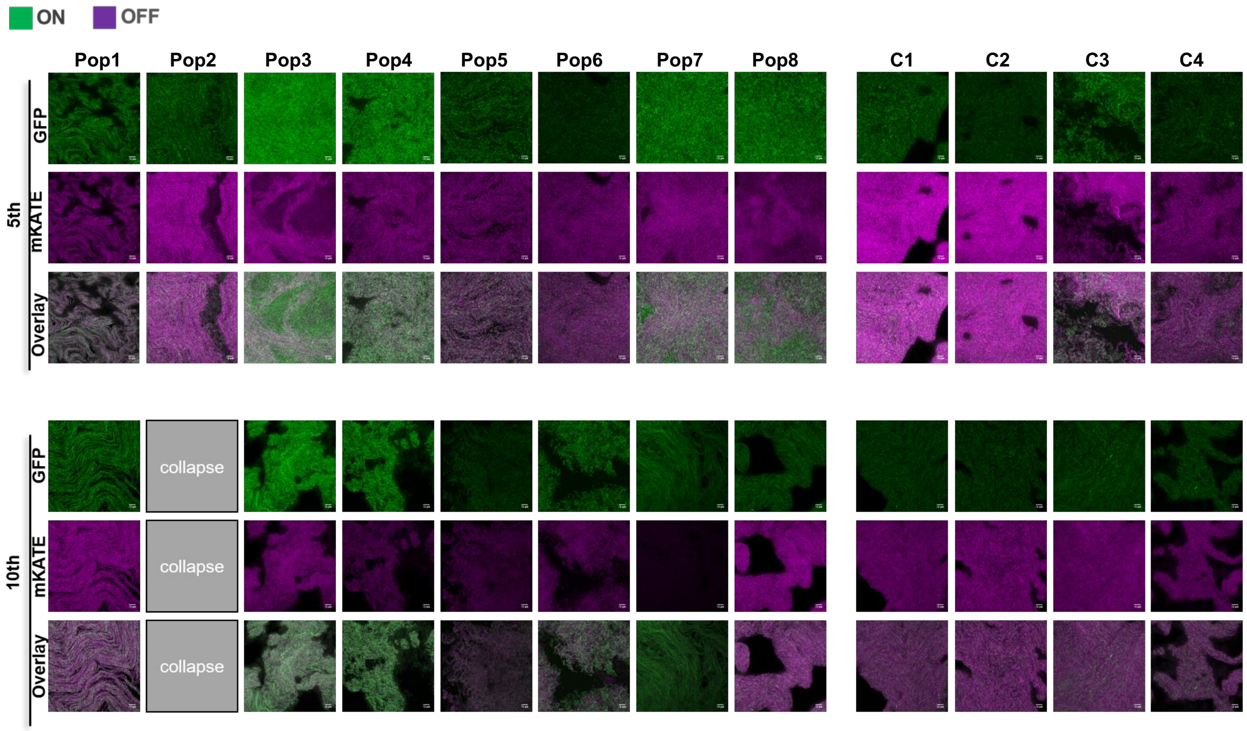


**Fig S2. Qualitative assessment of *eps* gene expression based on confocal laser scanning microscopy.** Pellicles formed by randomly selected WT strains 168 mKATE $P_{eps}$-GFP evolved in the presence of cheaters (Pop1-8) and in the absence of cheaters (C1-C4) were visualized using confocal laser scanning microscope. Cells constitutively expressing mKATE are represented in magenta (OFF cells) and *eps*- expressing cells (ON cells) are represented in green. Scale bar 10µm.

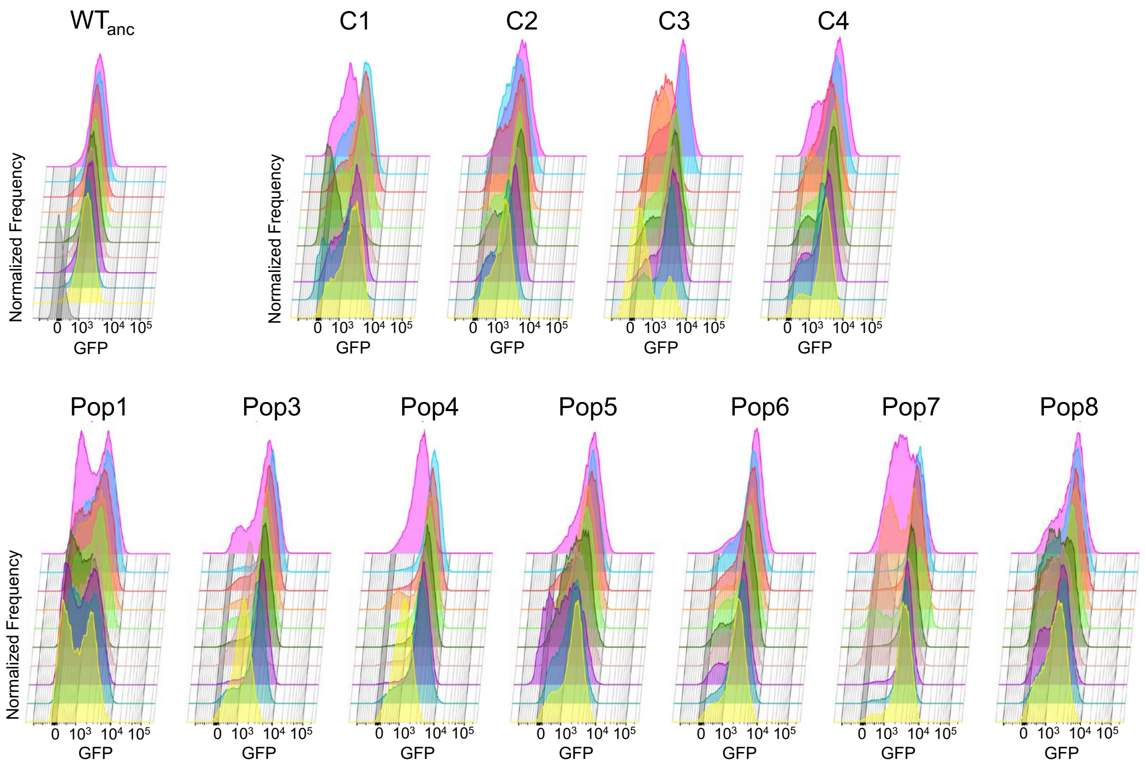


**Fig S3. Single cell level distribution of *eps* expression in 10 randomly selected single isolates from WT$_{anc}$ and evolved WT populations.** Flow low cytometry data (BD Facscanto II, BD biosciences) showing single cell level distributions of fluorescence intensity of 24-hour old pellicles established by 10 randomly selected single isolates from populations of WT evolved in the absence (C1-C4) or presence of cheaters (Pop1-8) as well as autofluorescence distribution in non-labelled WT control.

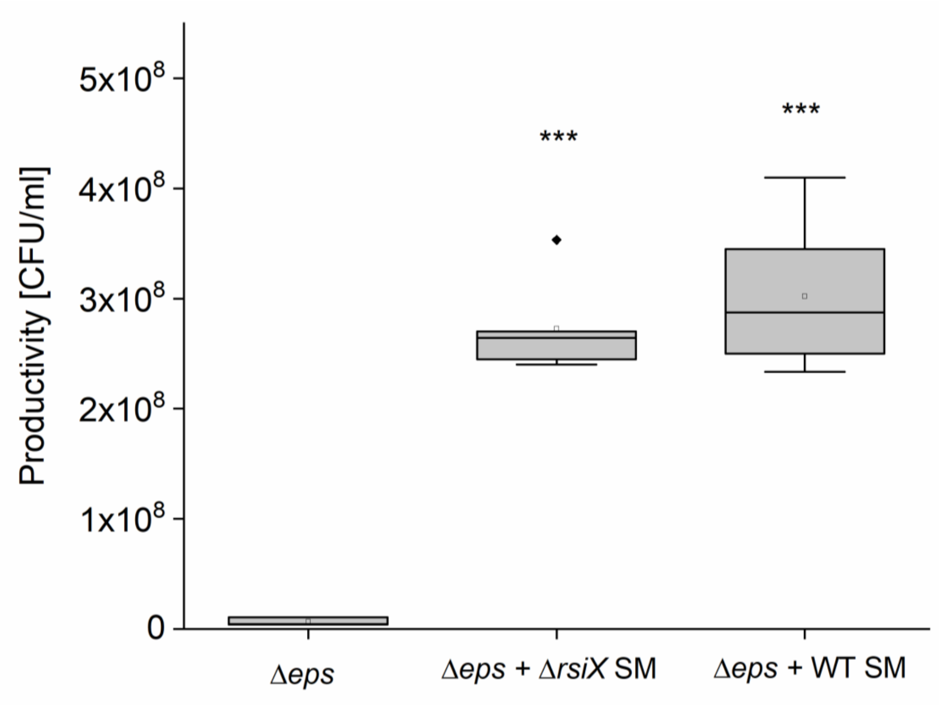


**Fig S4. Complementation assay of Δ*eps* with supernatant from Δ*rsiX* or WT.** Productivity data of pellicles produced by the complementation showed that hyper ON Δ*rsiX* mutant does not contribute to improved performance of Δ*eps*. Mean is represented in square inside the box plots; median is denoted by horizontal line within the boxes; whiskers represent the min and max (n=3 for Δ*eps*; n=6 for Δ*eps*+SMs). ***$p<0.001$ compared to Δ*eps*; Δ*eps*+ WT SM is not significantly different from Δ*eps*+ Δ*rsiX* SM ($p=0.58$) (ANOVA, Tukey Test).

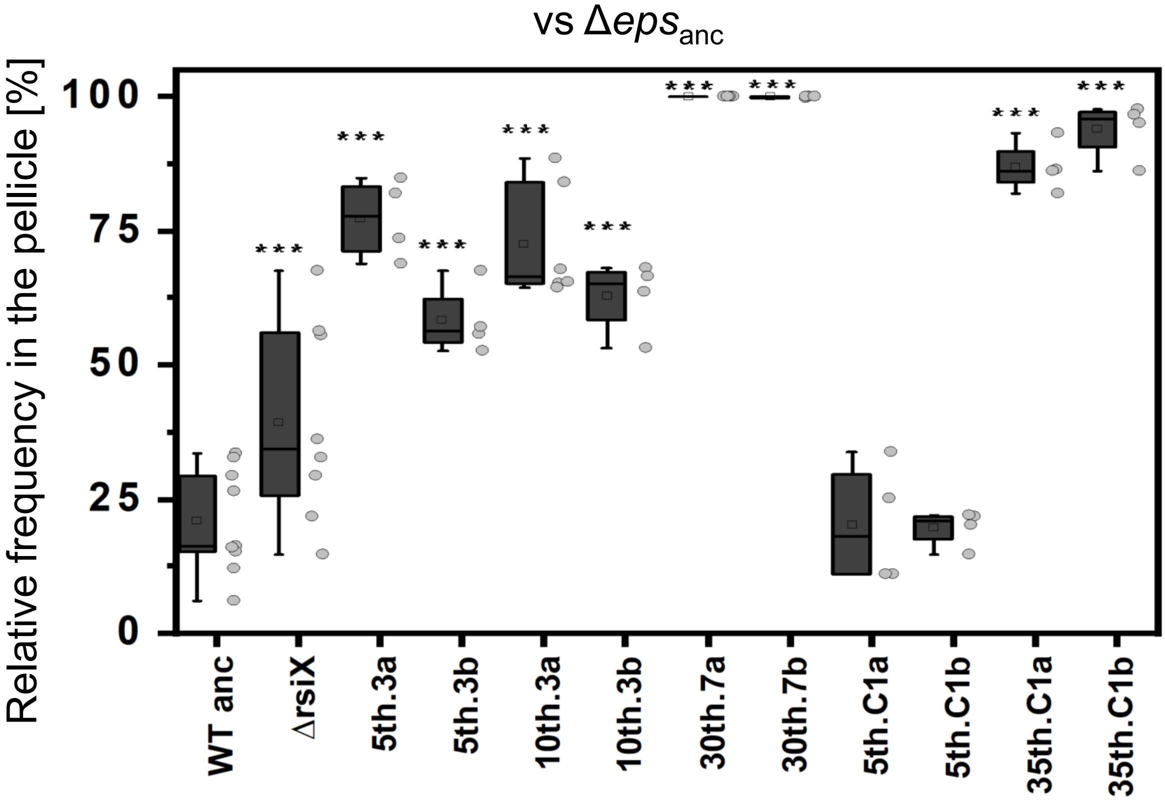


**Fig S5. Performance of Δ*rsiX* and evolved and co-evolved WT in mixed pellicles with Δ*eps*$_{anc}$.** Pellicle competition assay of single clones belonging to producer populations (WT$_{anc}$ (n=9), Δ*rsiX* (n=8), WT evolved with (n=4) and without cheaters (n=4)) against Δ*eps*$_{anc}$. Mean is represented in square inside the box plots; median is denoted by horizontal line within the boxes; whiskers represent the min and max; single dots represent the individual data points (n). ***p<0.001 compared to the WT$_{anc}$ (One-way Repeated Measures ANOVA, Dunnett Test).

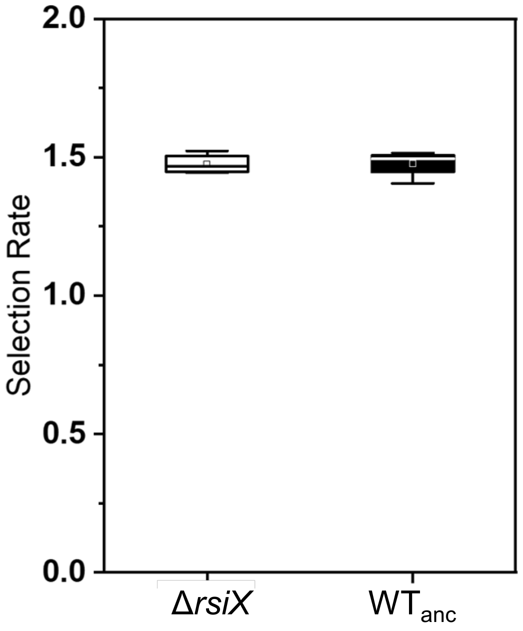


**Fig S6**. **Fitness effects of *rsiX* deletion.** Selection rate based on fitness assay in pairwise competition of Δ*rsiX* and WT ancestor showing no significant fitness cost brought about by *rsiX* mutation. Relative fitness of Δ*rsiX* is 1.00 ± 0.024 SD. Mean is represented in square within the boxplots; median is denoted by horizontal line inside the boxes; whiskers represent the min and max.

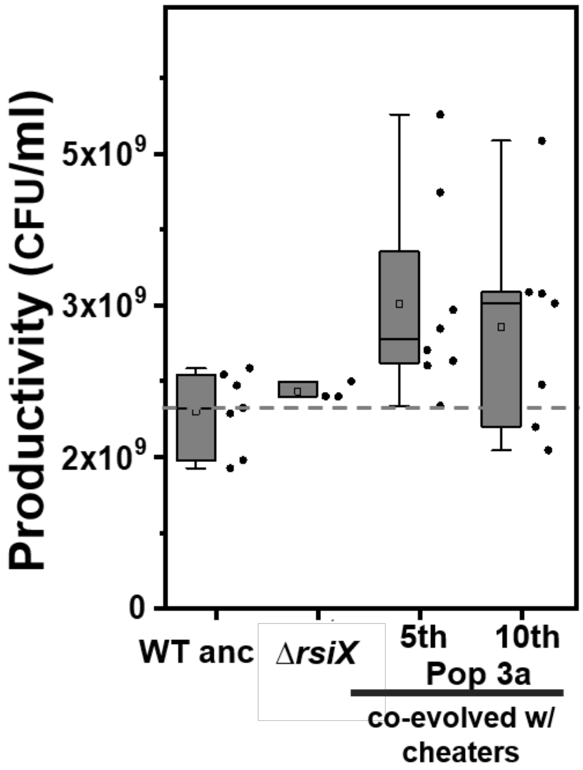


**Fig S7. Pellicle productivity of monocultures.** Total CFU/ml of pellicles produced by mono-cultures of $WT_{anc}$ (n=7), Δ*rsiX* (n=3) and evolved with cheaters (n=8) from population 3 ($5^{th}$, $10^{th)}$ and 7 ($30^{th}$) and single clones of WT evolved without cheaters (n=4). Mean is represented in square; median is denoted by horizontal line inside the box; whiskers represent the min and max; single dots represent the individual datapoints (n). All p values >0.05 (ANOVA, Tukey Test).

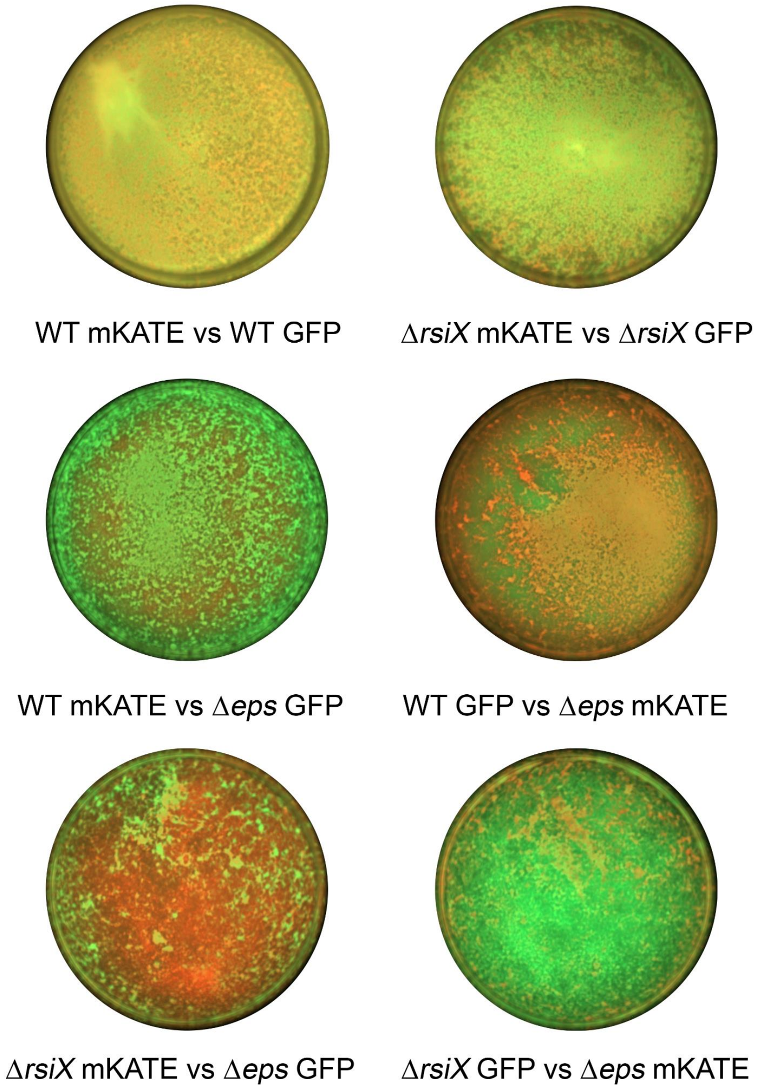


**Fig S8**. **Effect of *rsiX* mutation on positioning in the pellicle.** Competition assay between fluorescently WT+Δ*eps* and Δ*rsiX*+Δ*eps*. Strains labelled with constitutively expressed GFP and mKate proteins, were inoculated in 1:1 initial frequencies, pellicles were cultivated for 48h at 30$^{o}$C and visualized using stereomicroscope. Upper panels represent controls (two isogenic WT or ΔrsiX strains labelled with different fluorescent markers), middle panel represents pellicles formed by WT+Δ*eps* and bottom panels represent pellicles formed by Δ*rsiX*+Δ*eps,* each in two alternative combinations of fluorescent markers. Well size = 1.5cm.

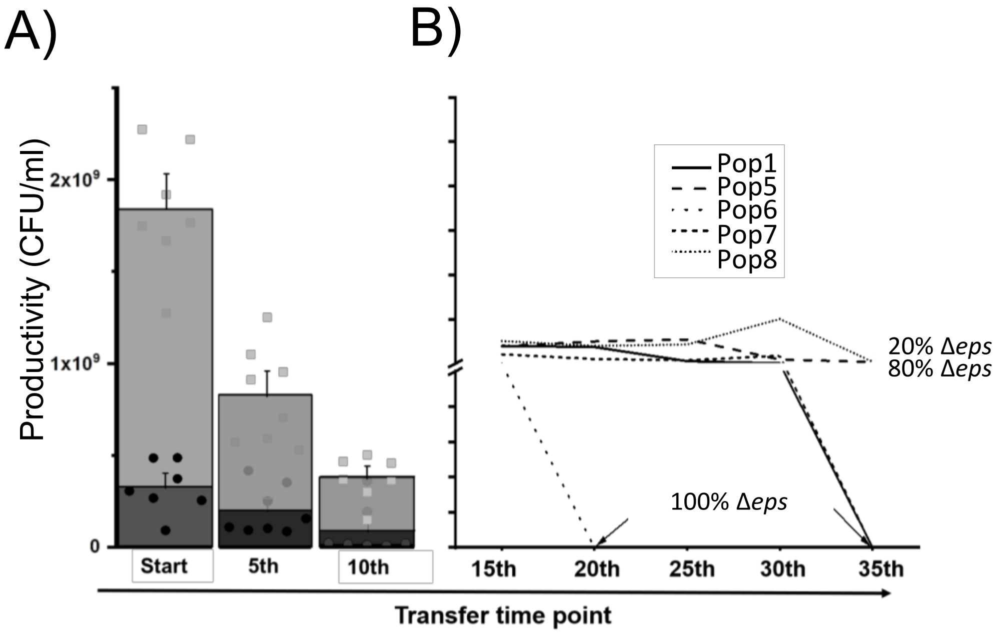


**Figure S9. Productivity changes during short and long-term co-evolution of WT+Δ*eps*. Pellicle** Total colony forming unit per ml of WT and Δ*eps* in 48-hour old pellicles **A)** non-evolved (start) (n=9), after experimental evolution at 5$^{th}$ (n=8 populations) and 10$^{th}$ transfer (n=7 populations) One population after 10$^{th}$ transfer was incapable to form pellicle attributed to WT being outnumbered by Δ*eps*. **B)** Line graph showing the fate of populations. Data was obtained from CFU assay using selective antibiotic marker, Kanamycin selecting for WT and Tetracycline for selecting Δ*eps.*
